# Analysis of Ultrafast Transient Absorption Spectroscopy Data using Integrative Data Analysis Platform: 1. Data Processing, Fitting, and Model Selection

**DOI:** 10.1101/165498

**Authors:** Evgenii L. Kovrigin

## Abstract

This manuscript describes a workflow for analysis of transient absorption (TA) spectroscopy data using Integrative Data Analysis Platforms (IDAP) software package. Time-dependent spectral series are analyzed through evaluation of the isosbestic point and kinetics of excited state and ground-state bleach decays. Model fitting and selection based on Akaike’s Information Criterion is discussed. As a practical example, we analyze excitation decays of a common protein label, Alexa Fluor 647.

---

Ultrafast transient-absorption (TA) pump-probe spectroscopy allows to monitor excitation-relaxation processes in chromophores with unmatched resolution^*1-4*^. In this method, the absorbance of the sample is measured before and after excitation (“pump”) pulse. The chromophore molecules in the excited state, typically, have a different absorption spectrum relatively to the ground state. By recording the sample absorption with a variable delay after the pump pulse one can observe a time-dependent spectral evolution that reflects relaxation of excited states back to the ground state. A number of commercial software packages are available for detailed analysis of the TA data such as *Surface Xplorer* (Ultrafast Systems LLC). We are developing the *Integrative Data Analysis Platform* (*IDAP*) as an open-source software package with a free academic license available from *http://lineshapekin.net/#IDAP*. The *IDAP* consists of a fitting engine with a generalized data interface for easy introduction of user-defined models, simultaneous analysis of multiple types of data, and statistical hypothesis testing. To allow for a customizable processing and analysis of the TA data, we developed the TA module for the *IDAP*, which is described in this manuscript.

## Materials

Alexa Fluor™ 647 C2 maleimide (A647) was purchased from ThermoFisher (Cat.# A20347). The dye was reconstituted with DMSO to obtain the 10 mM standard stock solution. The TA sample was prepared by diluting the standard stock of A647 with 25 µM L-cysteine (to quench the maleimide group) to achieve 7.5 µM solution concentration. Figure 1 shows an absorbance spectrum of the sample.

TA measurements were performed as described previously^*5*^. Data were recorded for one hour with 50 µW pump power. The duration of the pump and probe pulses were ca. 200 fs. The optical cell path length was 2 mm.

**Figure 1.**
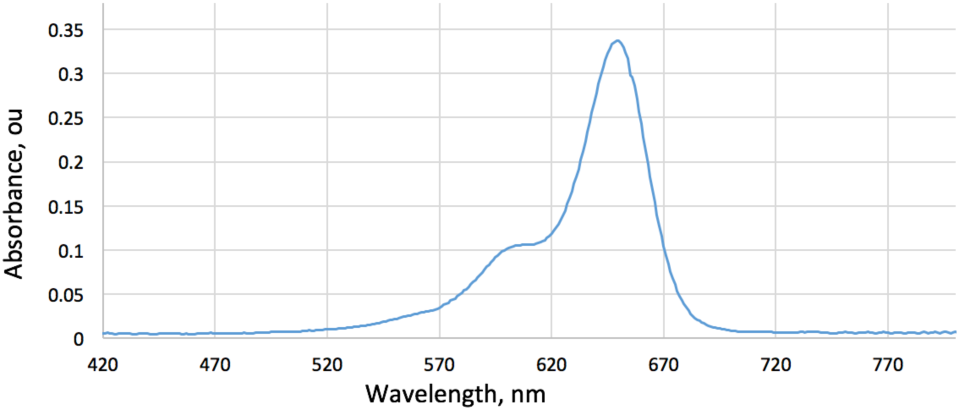
Absorbance spectrum of the A647 solution in a 2 mm optical cell.

## IDAP software

The *Integrative Data Analysis Platform* is written in MATLAB object-oriented M-code language. The TA data analysis is performed using a specialized data class *Exponential_Decay*, which includes a range of different multi-exponential models. The TA data are trimmed to start at 1 ps, therefore, a need for deconvolution of the Instrument Response Function (IRF) is removed. Some of the theory behind the analysis of the TA data with *IDAP* to extract quantitative FRET values and donor-acceptor distance distributions has been described in our earlier publication^*5*^. Here, we will focus on details of processing of the raw TA data with *IDAP* and multi-exponential model fitting and hypothesis testing. The TA control scripts for *IDAP* are included in Supporting Information.

## Pre-processing of the raw TA data

The two-dimensional TA data are exported from the TA acquisition software (in this paper: from the *Surface Xplorer*) to the comma-separated-values (CSV) format. The Python program *TA_convert_csv.py* prepares the datasets for import by IDAP (issue *TA_convert_csv.py ‘csv-file-name’*). The CSV file name is used to create a folder with extension *‘.data’* with two output files: *TA_data.txt* and *TA_parameters.txt*. The *TA_data.txt* is a rectangular matrix of intensity values with a row for each wavelength, and columns corresponding to different times. First row of the data matrix are the delay times. First column—wavelength values. The *TA_parameters.txt* includes the footer of the *Surface Xplorer* output with the instrument parameters (also displayed in the terminal window for inspection by the user).

## Data display and extraction of time-dependent decays

Visualization of the TA data and extraction of spectral regions of interest for analysis is performed with the control script *prepare_TA_data_main.m*. The *sample* variable is assigned in the beginning of the script to give a unique name to all figures and data objects generated in the session. Separate sets of settings are defined for different tasks. Uncommenting a selected set will direct *IDAP* to process the TA data in a desired way and deposit the results in a dedicated folder. The script generates a series of images: a small PNG file (suitable for email or web), a large PNG file (suitable for Powerpoint presentations), and a FIG file in MATLAB format (editable, exportable to EPS for publication). The extracted TA data are saved in the *session.mat* file for next steps of analysis (plotting and fitting).

### Overall view

Uncommenting “Overall view” section of the *prepare_TA_data_main.m* (with the rest of sections commented out) directs the *prepare_TA_data_main* to produce the view of a full two-dimensional dataset in two forms: a color map and a spectral overlay.

Figure 2 demonstrates the two-dimensional difference spectrum in the form of a color map with the time on X axis in a logarithmic format and the Y axis representing the absorption spectral range. The *lambda_min* and *lambda_max* variables set the display range for the Y axis. The red contours represent positive difference absorption values of the newly formed excited states while the blue contours indicate the ground-state bleach (GSB) due to the excitation pulse at 600 nm. The figure version that is saved in FIG format may be further adjusted using MATLAB Figure Editor to change color scheme and contour levels (this rule applies to all figures generated by *IDAP*).

**Figure 2.**
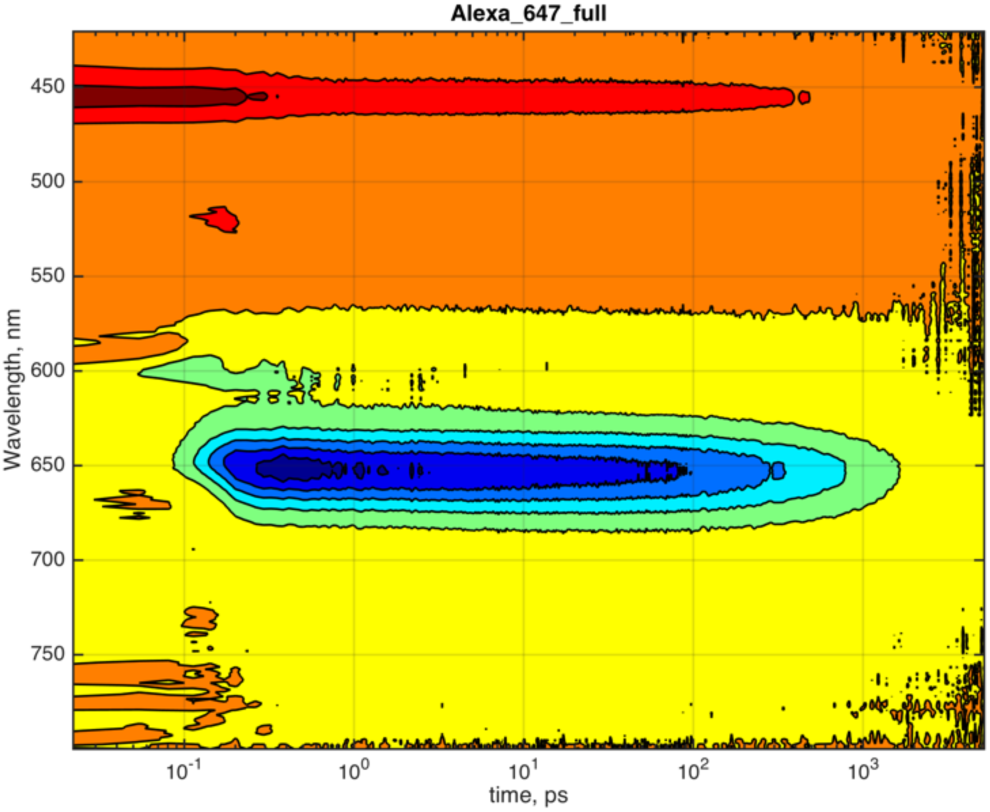
Two-dimensional color map of a transient absorption decay. Positive peaks are shaded red; negative—blue.

A series of slices are taken along the spectral dimension of Figure 2 at different delay times is shown in Figure 3. Each slice is obtained by integration of a specific time range such as illustrated in Figure 4. The desired time ranges are set by the *time_slices* variable in the beginning section of the *prepare_TA_data_main.m*.

**Figure 3.**
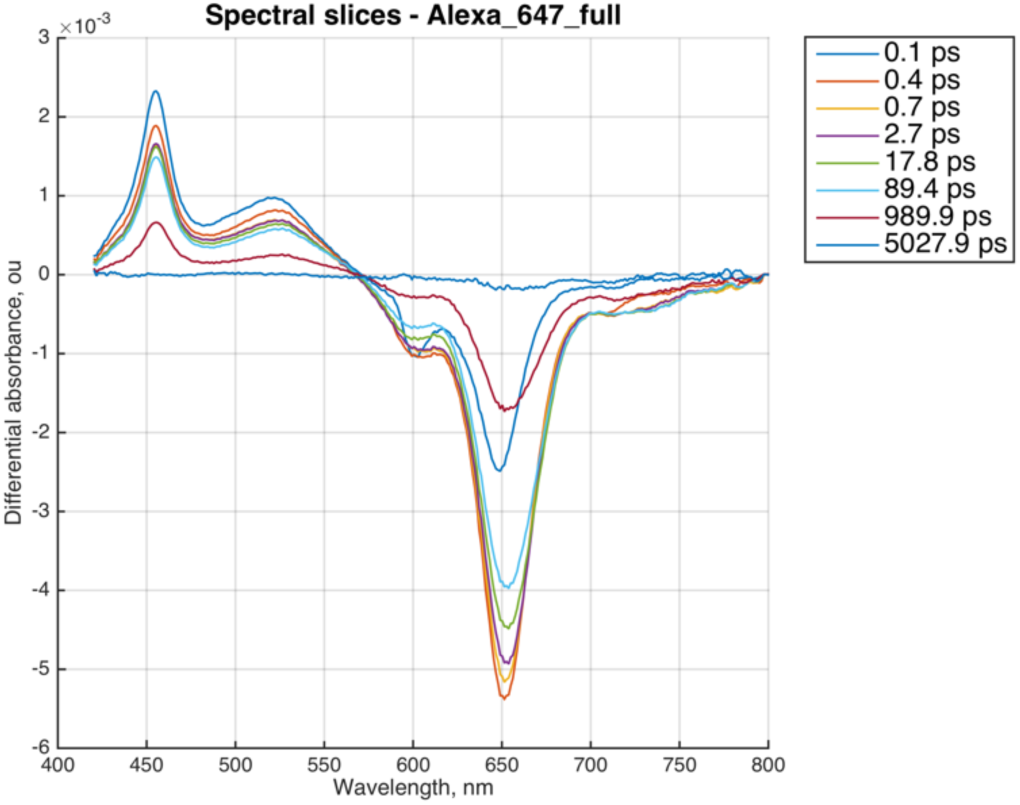
Spectral slices along the Y axis in two-dimensional dataset taken at different time intervals between the pump and the probe pulses.

**Figure 4.**
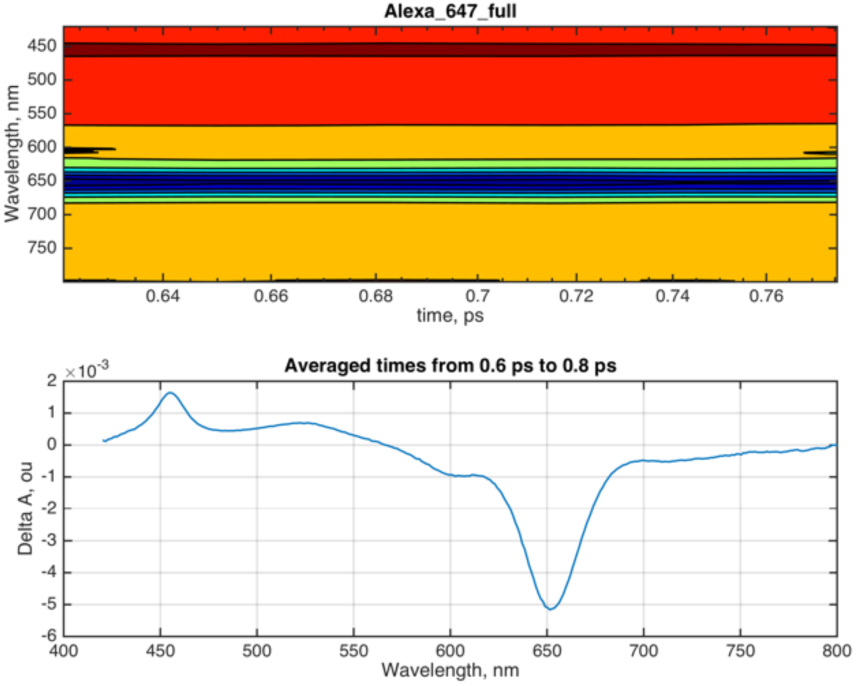
Example of the two-dimensional spectral region (top panel) extracted for integration to obtain the spectral slice corresponding to 0.7 ps (bottom panel).

### Isosbestic point

If the major excitation-relaxation mechanism is a transition to a single excited state relaxing directly into the ground state (a two-state transition) the spectral overlay such as in Figure 3 reveals an isosbestic point—a wavelength where spectral intensity remains constant during entire process. Figure 5, top panel shows a zoomed-in view of the isosbestic point observed near 570 nm (uncomment “Zoom on isosbestic point” section and rerun the *prepare_TA_data_main.m*). Figure 5, bottom panel shows time evolution of the isosbestic point region. The *slices* variable defines a (narrow) spectral region to calculate the average spectral intensity for each time point. Initial perturbation in this spectral range is visible during the excitation laser pulse within the first 0.2 ps.

**Figure 5.**
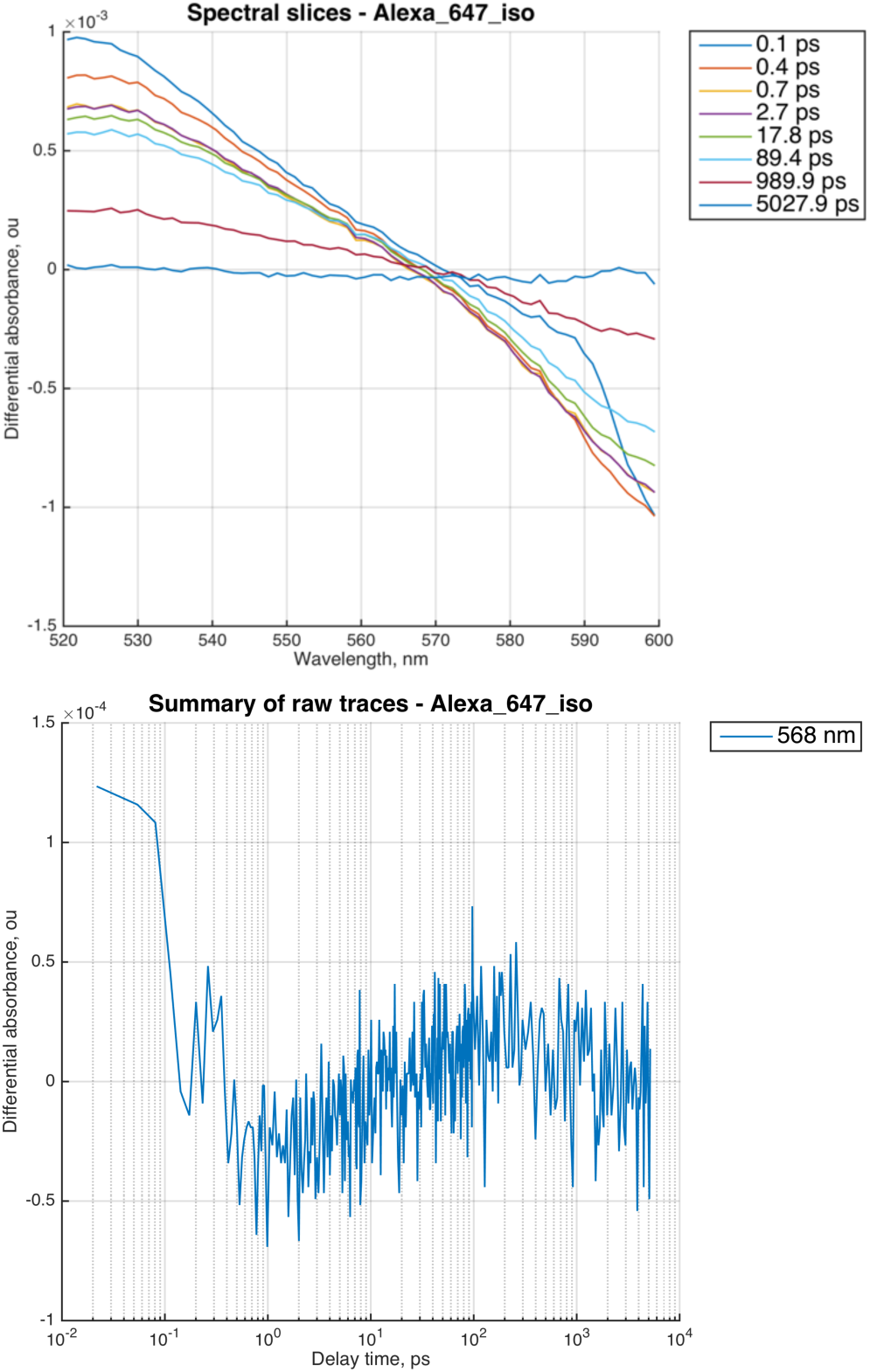
(Top) Expanded spectral area near the isosbestic point. (Bottom) Time-dependent spectral intensity integrated from 566 nm to 570 nm for each delay time.

### Excited state region

Excited states of the chromophore created by the pump pulse at 600 nm are observed as positive peaks in Figure 2 and Figure 3. For a more detailed look, the section “Zoom on positive peaks” expands this spectral region (Figure 6) and generates the time-dependent traces at the user-defined wavelengths (Figure 7). The wavelength labels correspond to a middle of the spectral ranges in the *slices* array. In Figure 7, each time point is an average intensity in the 450-460 nm and 520-530 nm ranges, respectively. One can see initial perturbation of the traces when excited states are created by the pump pulse followed by a decay of the excited state absorbance. The *signal_sense* variable is set to +1 to indicate to *IDAP* that we are watching the positive peak decay (as opposed to negative features). In principle, this determination might be made automatically based on the signal intensity. However, with weak signals, the particularly noisy data points may have opposite signs throwing off the automatic estimate. Therefore, the peak “sense” is enforced by the user directly.

**Figure 6.**
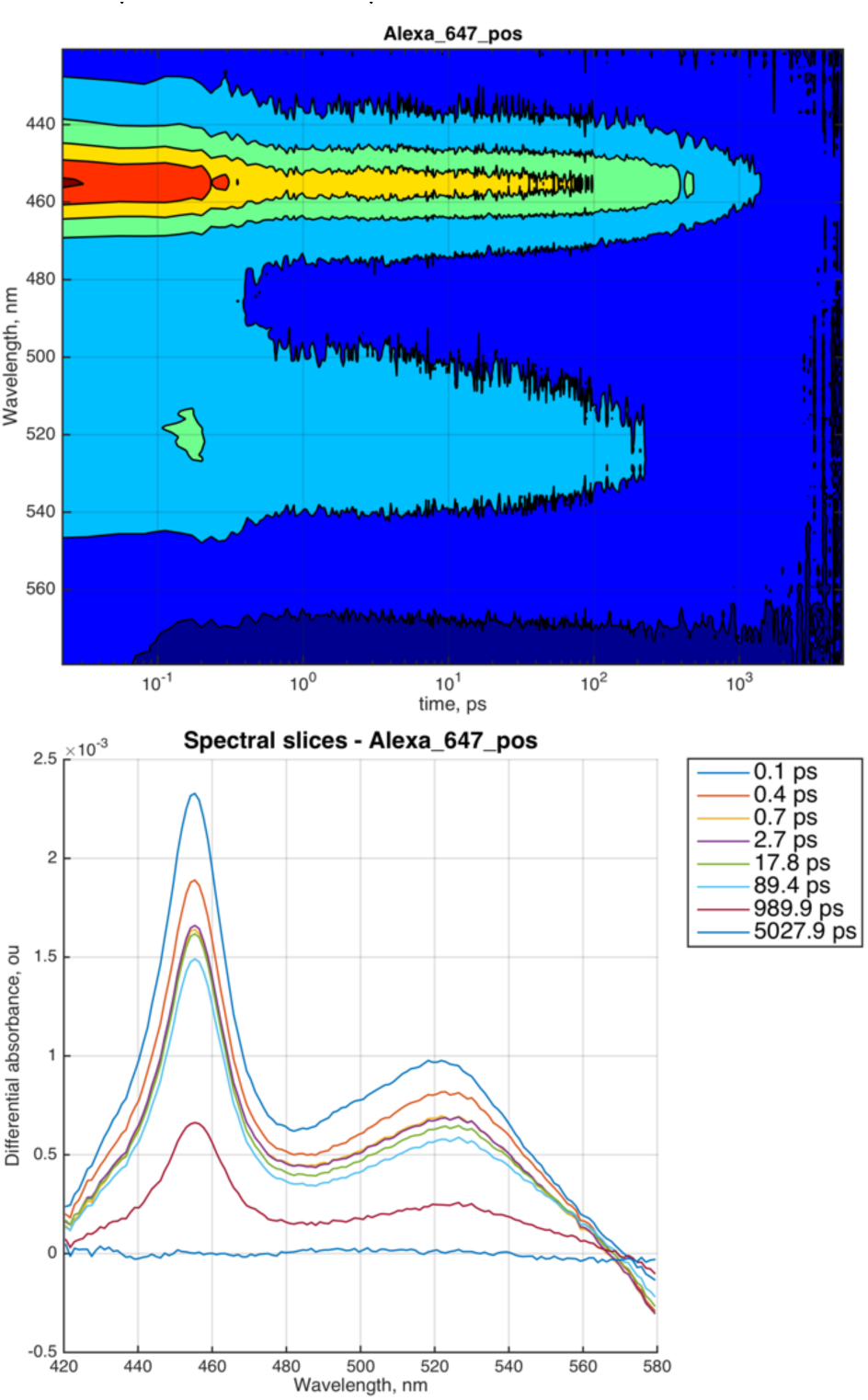
Excited state region of A647 TA dataset. (Top) Two-dimensional colormap of the dataset. (Bottom) Spectral slices in the excited state regions.

**Figure 7.**
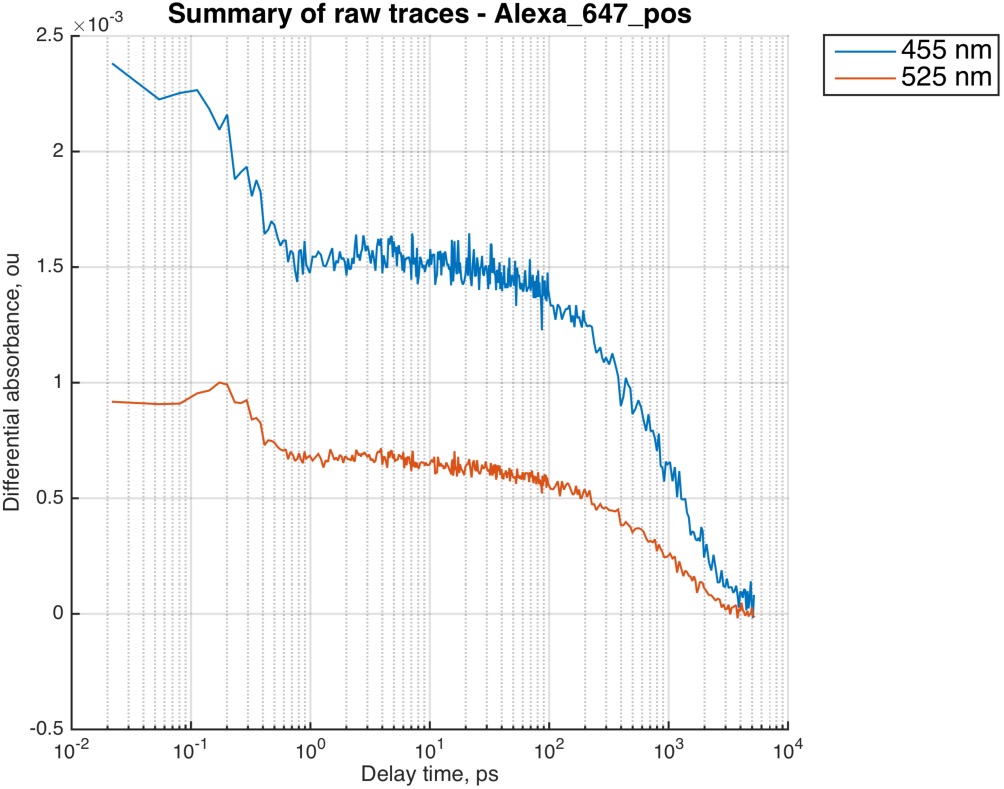
Time-dependent decay of the differential absorbance of the excited states at chosen wavelengths.

For the accurate extraction of time-dependent traces, it may be desirable that the baseline offset is subtracted from the data. This is controlled by the *offset_correction* variable (‘yes’/’no’), and the *X_start_baseline* and *X_end_baseline* parameters (common for all sections) defining the data region in the beginning of a trace prior to the pump pulse. Figure 8, top panel, shows examples of the baseline regions corresponding to the datasets in Figure 7. It is essential that the user inspects the baseline region to ensure that the *IDAP* calculated a reasonable baseline offset. The average value of the baseline in Figure 8, top panel, is subtracted from the raw absorbance values to the give baseline-corrected trace. In the next step, the dataset is trimmed to start at 1 ps (defined by *X_trim_from* variable) and normalized by the maximum point. Normalization is only done to facilitate comparison of traces with different absolute amplitudes and will not affect model fitting in the future steps. The resulting signals are displayed in Figure 8, bottom panel, smoothed to facilitate comparison. Negative orientation of the axis is arbitrarily chosen to match direction of GSB signals (next section) and will not affect fitting results.

**Figure 8.**
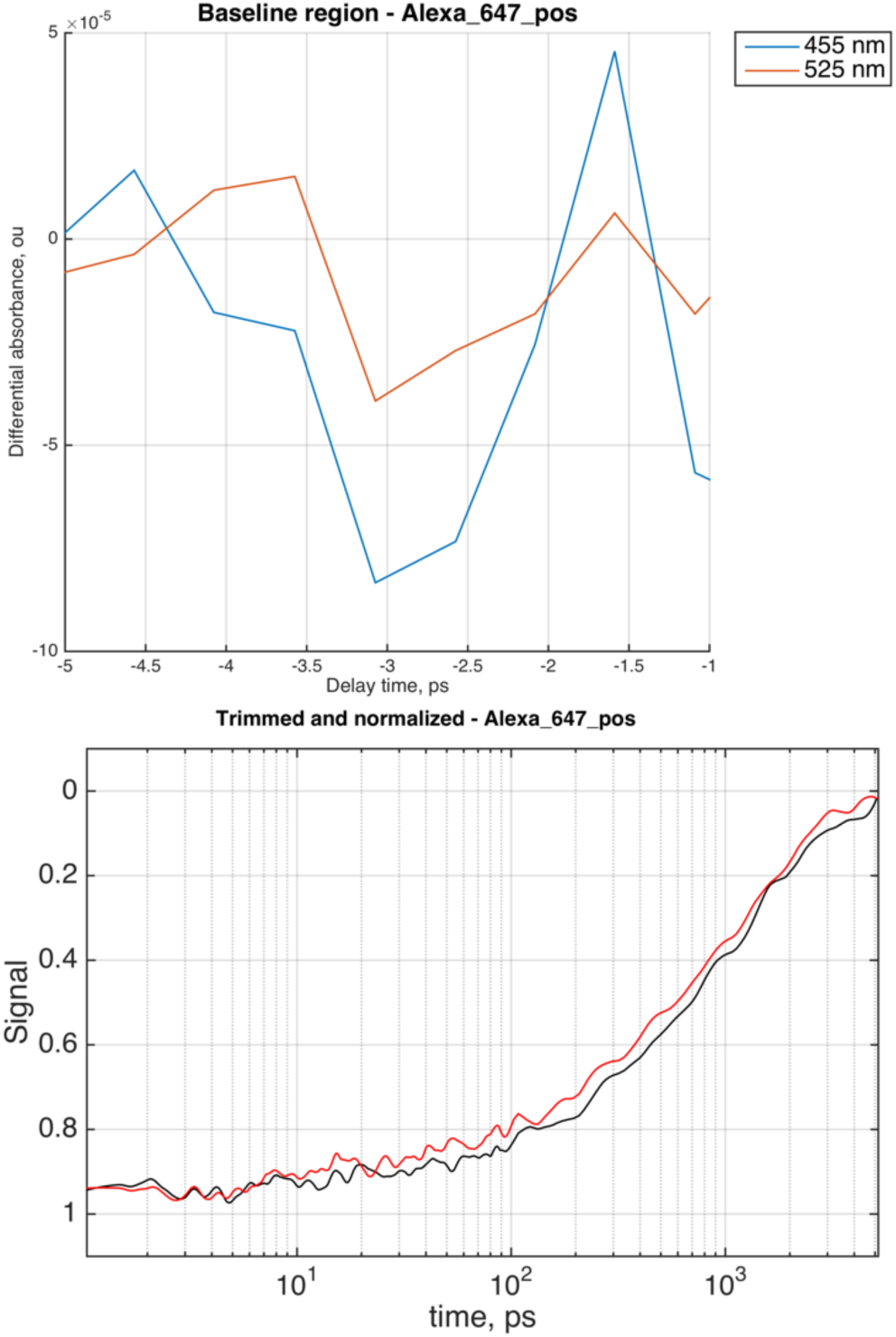
Preparation of the decay data at chosen wavelengths for further analysis. (Top) Time range displaying the preexcitation baseline. (Bottom) Trimmed, baseline-corrected and normalized decays. Black trace, 455 nm; red trace, 525 nm. The data is smoothed with the moving average window for plotting.

### Ground state bleach

Figure 9 visualizes the negative peak of the ground-state bleach of A647 (uncomment the “Zoom on negative peaks” section). The ground-state population of the chromophore molecules was reduced by the excitation process, therefore, there is less absorbance in the area where the steady-state absorption maximum is (compare the bottom panel of Figure 9 with Figure 1). Extraction of time-dependent decay traces is done similarly to the excited state decays with the only difference that the *signal_sense* variable is set to -1.

**Figure 9.**
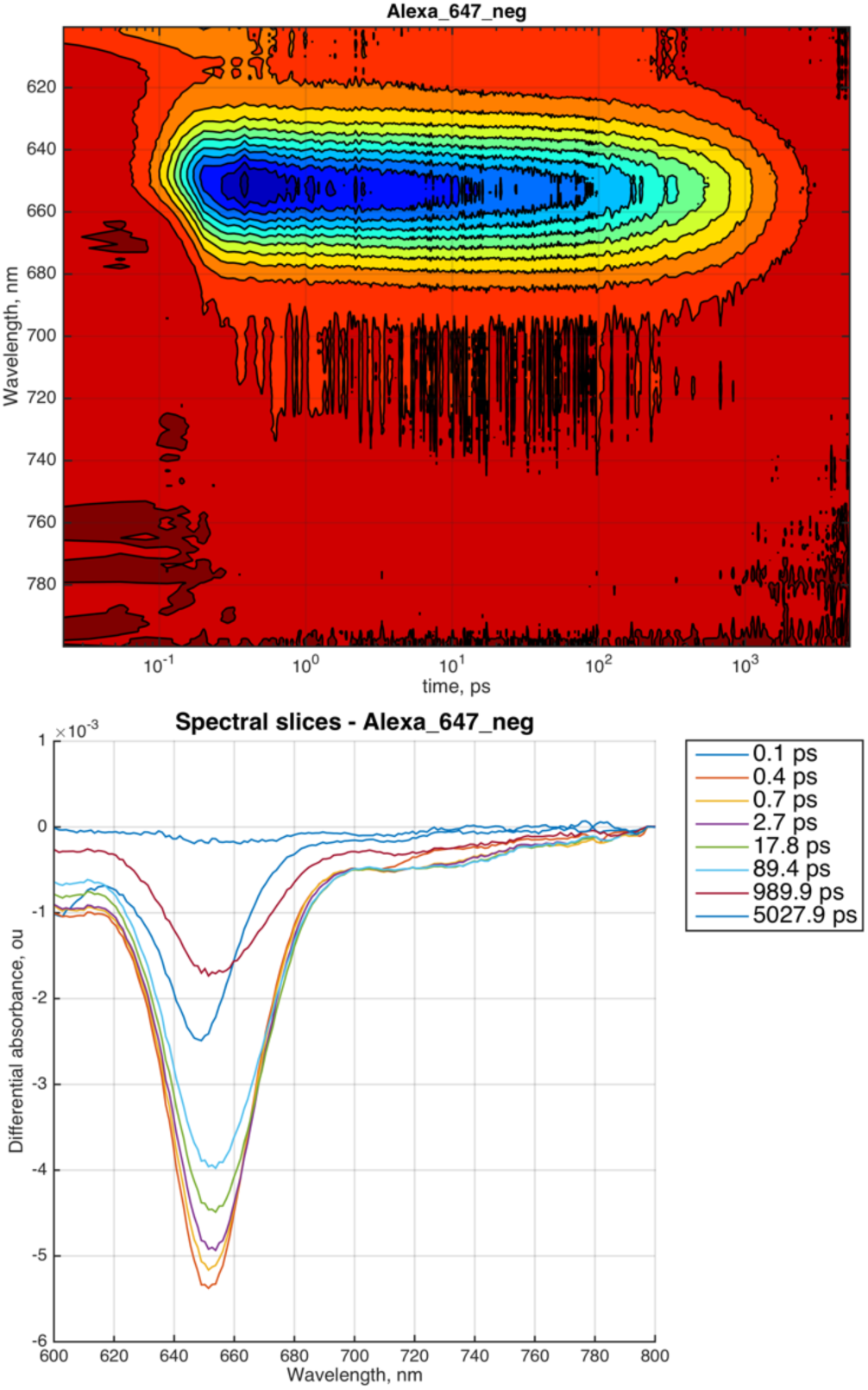
Ground-state bleach of the A647. (Top) Color map of the dataset; (Bottom) Overlaid spectral traces at different delay times.

To examine the time-dependent GSB decay, we selected the absorption maximum region (645 to 665 nm) and also a spectral range on a shoulder of the absorption peak around 720 nm. Top panel of Figure 10 shows the raw decays reflecting expected difference in the their amplitudes. The bottom panel shows trimmed and normalized data graphed with a moving average window to facilitate comparisons. The kinetics of the two GSB traces appear different, yet considering very low signal-to-noise ratio (S/N) of the 720 nm trace, one should refrain from making conclusions from this dataset. For a proper comparison, an additional experiment is required with longer acquisition and/or higher concentrations of the dye to bring S/N in the 720 nm decay down to a level comparable to the S/N of the 655 nm trace in Figure 10.

**Figure 10.**
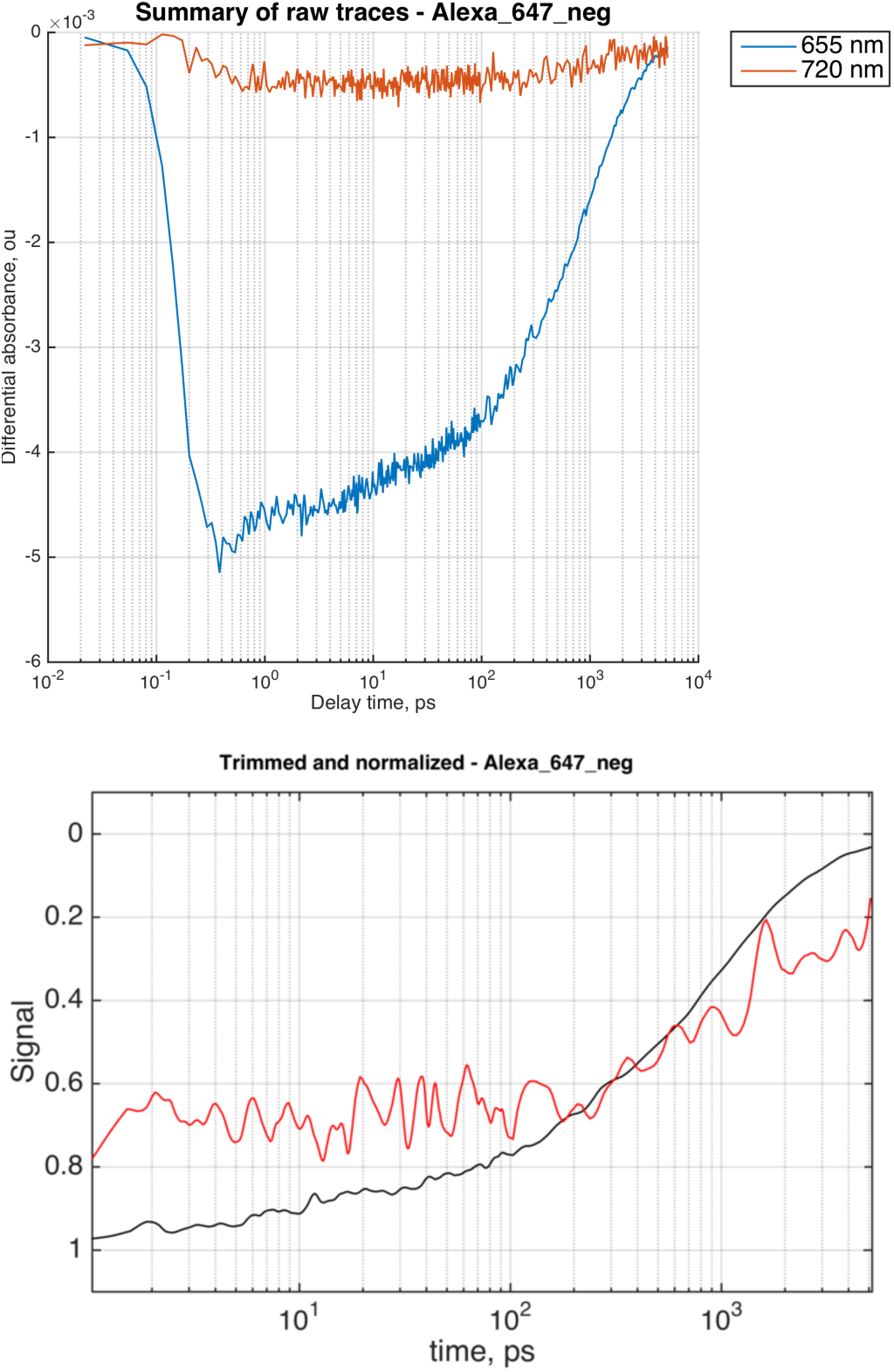
Ground-state bleach of the A647. (Top) The raw decay data at two wavelengths. (Bottom) Trimmed, baseline-corrected and normalized decays. Black trace, 655 nm; red trace, 720 nm. Normalization is formally performed using a maximum point of the raw data before smoothing—explaining offset of the two traces from 1. Smoothing is only done to generate the trace for display and will not alter the data for fitting.

### Comparison plotting

Multiple time-resolved traces originating from the same or different TA experiments may be plotted together with the *plot_multiple.m* script. The *datasets_files* is a cell array with full (or relative) paths to the *session.mat* files of the corresponding datasets. Figure 11 demonstrates overlay of all traces examined in the A647 dataset with GSB decays multiplied by -1 (the *signal_sense* value) for easier comparison.

**Figure 11.**
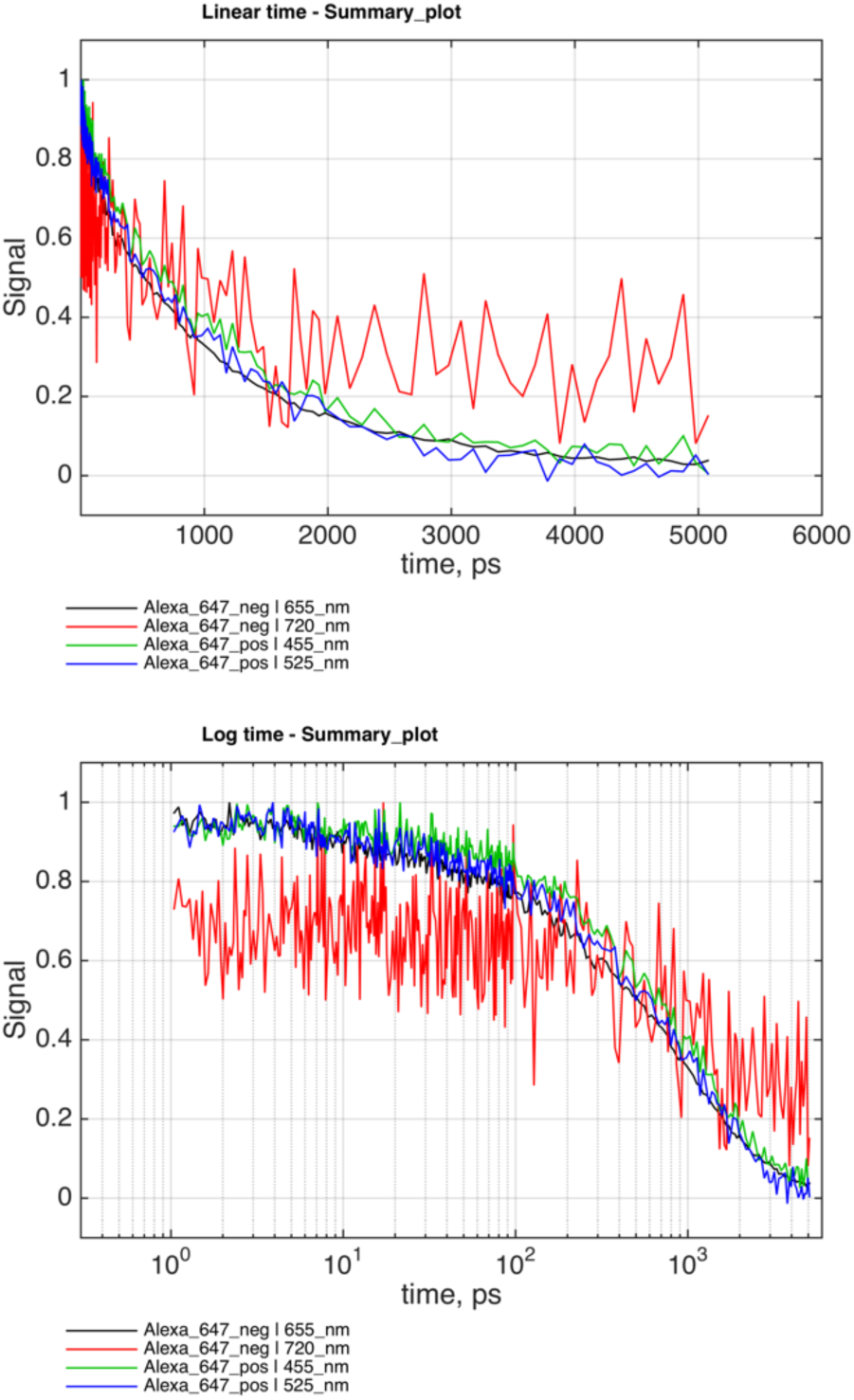
Summary of the time-dependent differential absorption traces with linear (top panel) and logarithmic (bottom panel) time axis. Solid line connects raw data points (no smoothing applied).

## Data fitting with alternative models

The *session.mat* files containing multiple decays obtained in the previous sections are directly usable in the next fitting phase of analysis. For purposes of easier data management, it is still preferable to re-run *prepare_TA_data_main.m* to extract chosen traces one at a time such that each *session.mat* contains only one trace (use sections “Zoom on one feature”). In the example below, we extracted a single GSB trace by integrating between 645 and 665 nm (corresponding to the “655-nm” trace in Figure 10).

The fitting process is controlled by *fit_exp_decay_main.m* script. The script may be run in three modes: (1) display of data and initial approximation of the model to the data without fitting, (2) least-squared fitting of the chosen model to the data, (3) mutliple runs with synthetic datasets to determine parameter uncertainties and correlations through the Monte-Carlo analysis. To request a specific mode, the *fit_mode* parameter is assigned the ‘show-data-only’, ‘fit-only’, and ‘Monte-Carlo’ values, respectively.

There is a choice of fitting algorithms including simplex and other algorithms supplied with the MATLAB Global Optimization toolbox. In our experience, the simplest algorithm ‘simplex’ was also the most effective. Parameters of fitting algorithms may be adjusted by editing their values in *IDAP* code (*TotalFit.m,* section “Options for fitting routines”).

Fitting model is set using the *current_model* variable to one of the multi-exponential models. These models are sums of exponentials with the individual amplitudes *a*_*i*_ normalized to unity and an adjustable overall amplitude *A*:

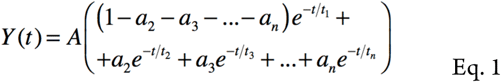

where *t*_*i*_ are time constants (lifetimes) of *N* individual exponential terms. Descriptions of the models and their parameters are given in the code of *Exponential_Decay.m* and listed in *‘+exponential_decay_laws/’* folder.

It is very important to provide reasonable initial parameters to the fitting routine such that the model approximates data close enough at the start of the parameter optimization. We start fitting with a single-exponential model (*current_model=‘norm-exp-1’* and *parameter_modes=[1 1]*) and estimate the first lifetime from the decay trace shape plotted with the logarithmic time axis such as Figure 10. The maximum slope of the curve for GSB at 655 nm is observed around 900 ps. Setting *t1=900* and *fit_mode=‘show-data-only’* produces a reasonable starting condition for a fit because it captures the major change in the data (Figure 12). The images of initial approximation plots are deposited in the folder with an extension *‘Initial’* including the sample name, the optimization algorithm, and the model name.

**Figure 12.**
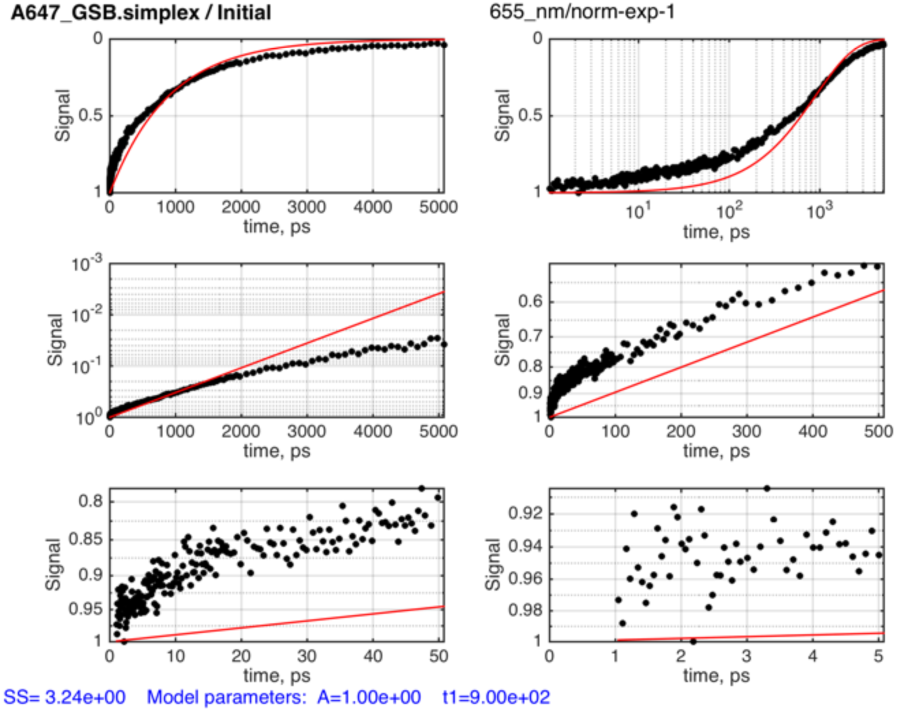
Examination of starting parameters for fitting A647 GSB at 655 nm with a single-exponential function. Raw data (black dots) and model (red line) plotted in linear amplitude and time coordinates (top left), logarithmic time (top right), linear time and logarithmic amplitude (middle and bottom rows) with different time windows.

The parameter optimization is initiated by uncommenting *fit_mode=‘fit-only’* and re-running the control script. The fit results are found in the folder with extension *‘Fit_Only’.* Figure 13 shows the optimization result. The parameters of the best fit are given in the bottom of the figure and in the *adjustable_parameters_only.txt* and *all_datasets.txt* files (the ‘*Best_Fit/’* folder). Note that the sum of squares (SS) value shown in the figure is the total SS for the dataset, while the text files contain SS normalized ‘per point’. It is clear that we need more than one lifetime to characterize this GSB decay. Inspection of the Figure 13 suggests that the lifetime of about 10-50 ps accounting for 20% of the total amplitude may also be needed to improve match to the data.

**Figure 13.**
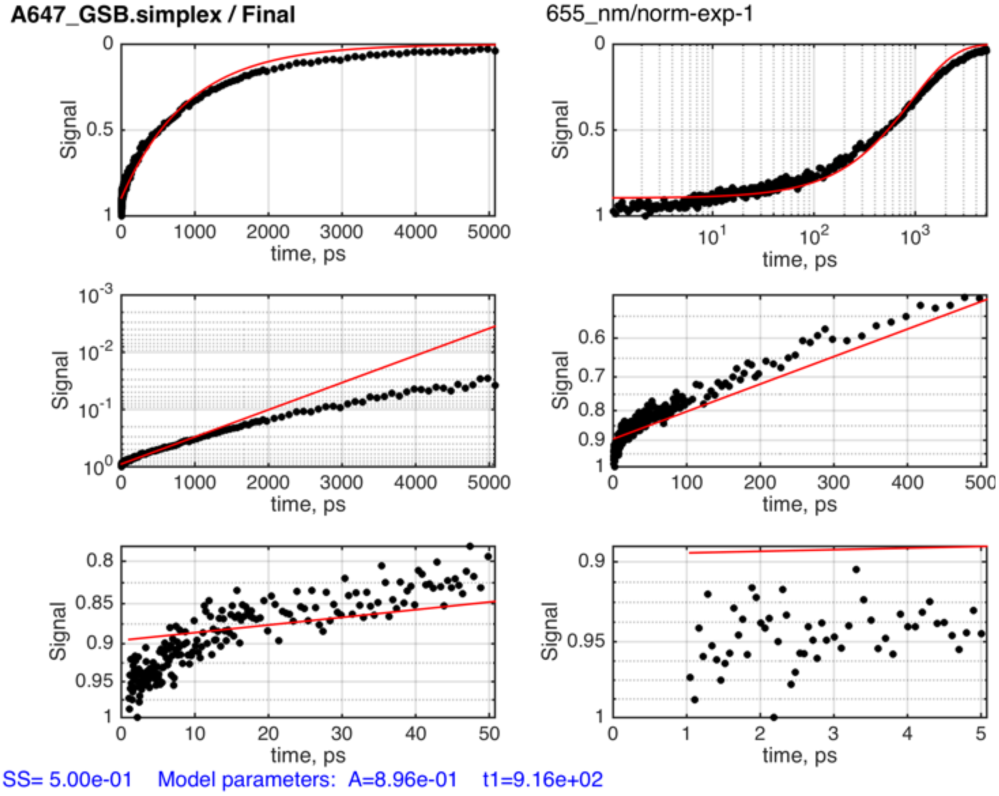
Fitting of A647 GSB with a single-exponential function.

When switching to the two-exponential model *‘norm-exp-2’* one also switches the value of *parameter_modes* to the array *[1 1 1 1]*. The modes are ‘1’ for any parameter to be treated as variable or ‘0’ for keeping it fixed to the set value. If one desires to change parameter modes, the order of parameters in the model may be looked up in the file *Best_Fit/all_datasets.txt* created after starting the script in ‘show-data-only’ mode. In this manuscript, we will always keep all parameters as adjustable.

Setting *a2=0.2* and *t2=10* results in a better fit shown in Figure 14. The SS is significantly reduced, yet the model curve deviates from the data at longer times. A “stem plot” with the lifetimes and their amplitudes allows for easy examination the best fit results (Figure 15). It is important that the starting parameters for every next stage are adjusted to the best-fit parameters of the previous optimization run.

**Figure 14.**
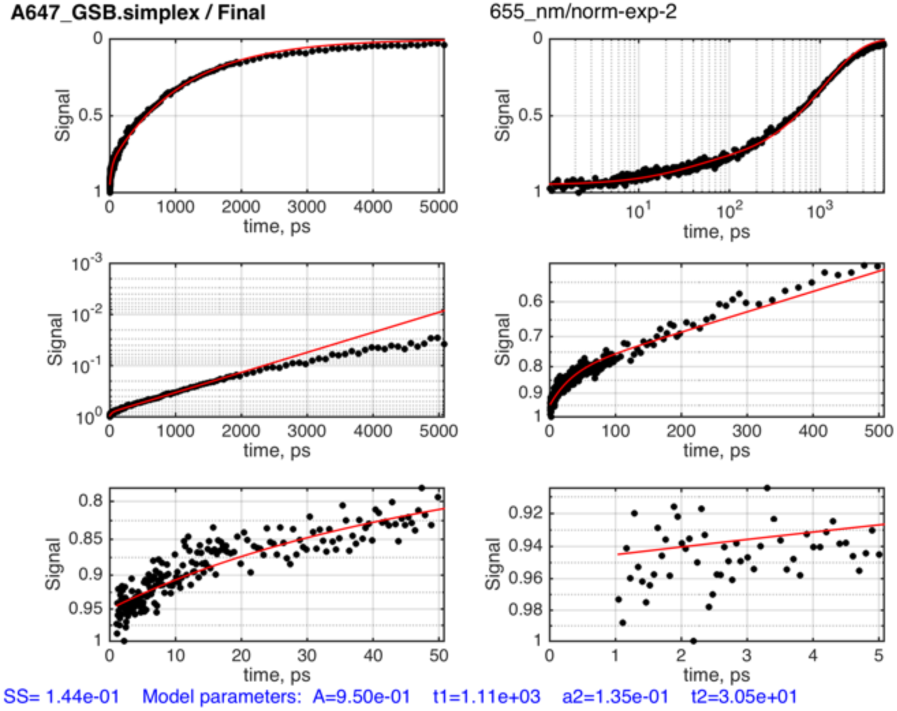
Fitting of A647 GSB with a two-exponential function.

**Figure 15.**
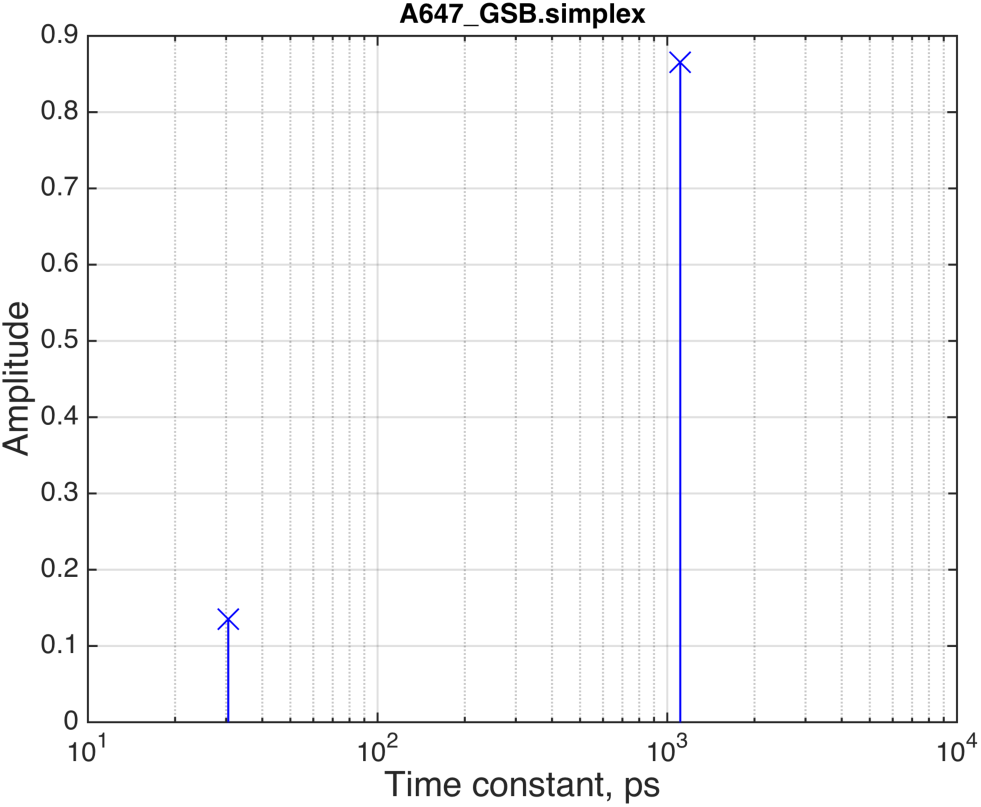
Optimized parameter values for the two-exponential function. Amplitude of the each exponential term is plotted versus its corresponding time constant.

A deviation of the model from the data is observed at longer times (Figure 14, middle left) while the difference in amplitude is quite small (Y axis is logarithmic in this pane). Addition of the third lifetime on the order of 4 ns and with relatively small amplitude (*t3=4000, a3=0.05*) leads to a shift of the lifetime distribution and a better fit (SS is reduced by 30%) shown in Figure 16. Notable, that our guess that a longer lifetime is what the model needs most proved incorrect. Instead, a lifetime of 300 ps was added improving the model match to the data in sub-nanosecond time range (Figure 17).

**Figure 16.**
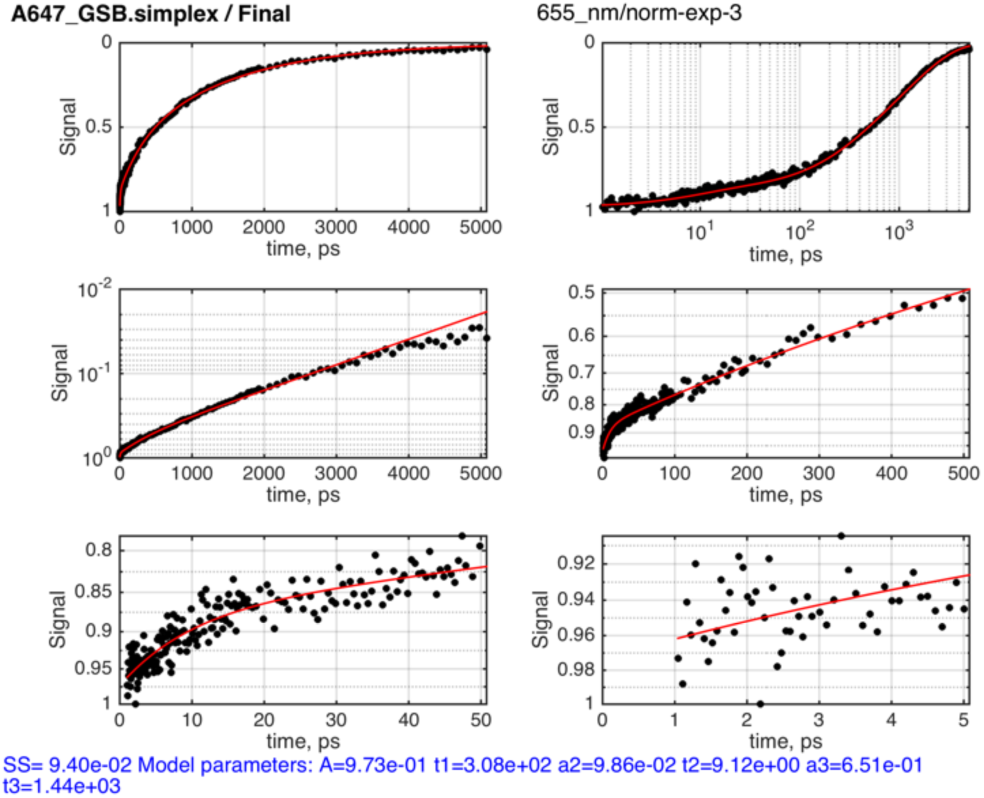
Three-exponential fit.

**Figure 17.**
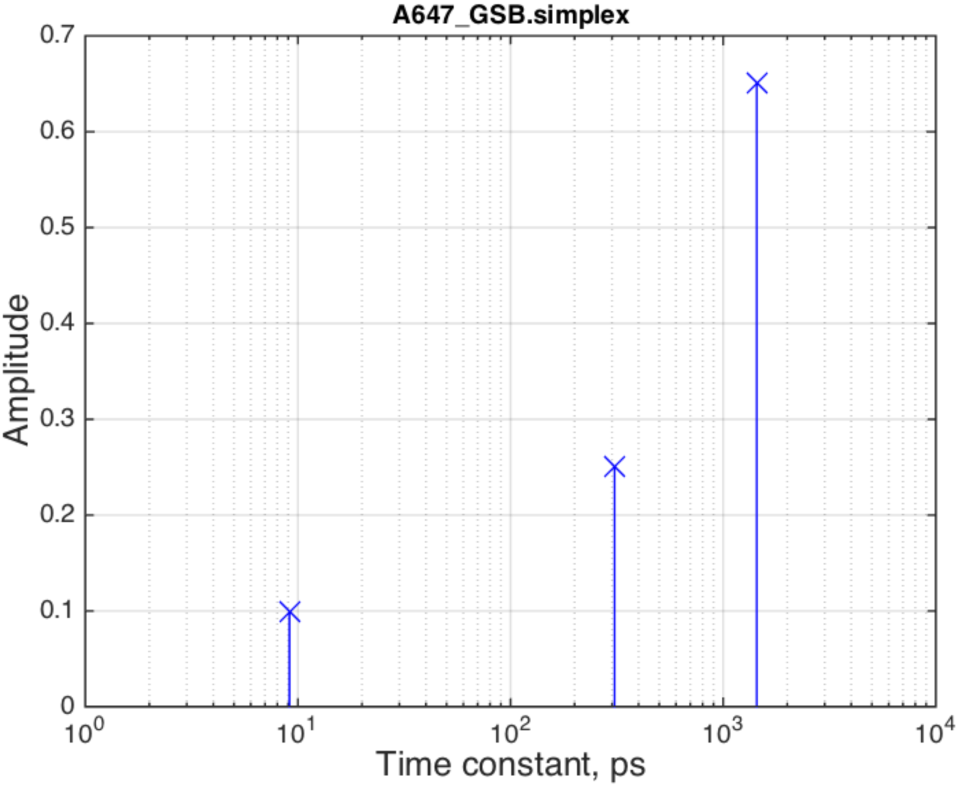
The optimized parameter values for the three-exponential function.

The middle-left pane of Figure 16 still indicates that some reduction of sum of squares may be gained by adding a very long lifetime with a very small amplitude. Setting *t4=10000, a3=0.01* produces further better fit shown in Figure 18. Thus, all features of the GSB decay appear to be accounted for by including four exponential terms in the model. It is notable that representation of the best-fit amplitudes vs life-times as in the bottom panel of Figure 18 may be directly correlated to appearance of the GSB decay in the logarithmic time scale (Figure 18, top panel, top right pane). The relative height of the “stems” visually correlates to fraction of GSB intensity change within the corresponding time range with t_3_ contributing most and t_4_ contributing least.

**Figure 18.**
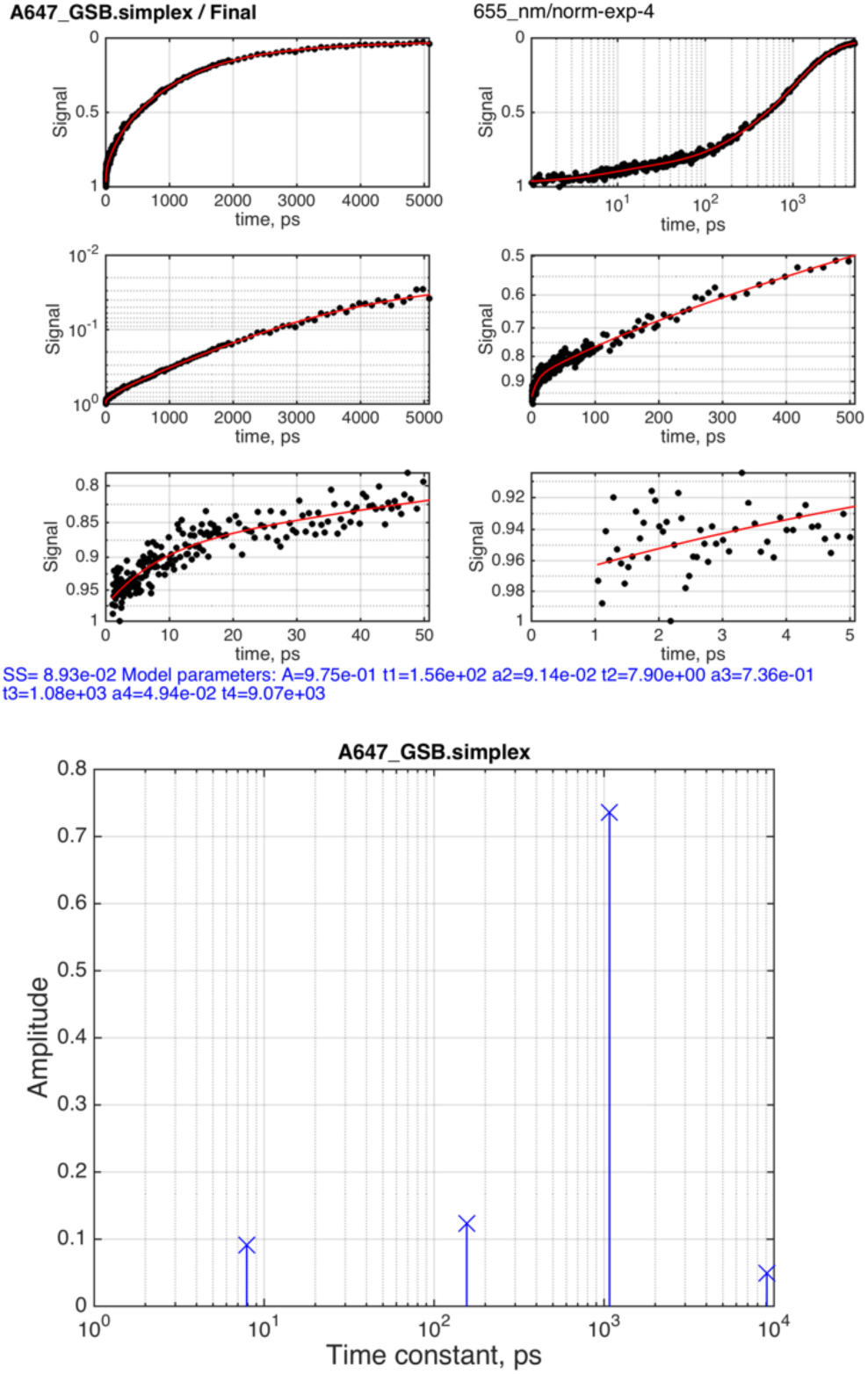
Four-exponential fit.

Further increasing number of exponential terms leads to further reduction of sum of squares (Figure 19) but involves a new term with a lifetime close to a picosecond. Processes this fast have to be analyzed in a more complex fashion using deconvolution of the Instrument Response Function (IRF) because this time scale is similar to the width of the pump pulse of 200 fs and time constants of the detector (implemented in *SurfaceXplorer*). The most important observation is that amplitude of this picosecond term is negative, indicating that it is a *rising* exponential component not decaying one. We may or may not consider such observation meaningful depending of what we know of the relaxation process in the chromophore. Addition of the sixth exponential terms does not bring further improvement and also includes a negative sub-picosecond lifetime (not shown).

**Figure 19.**
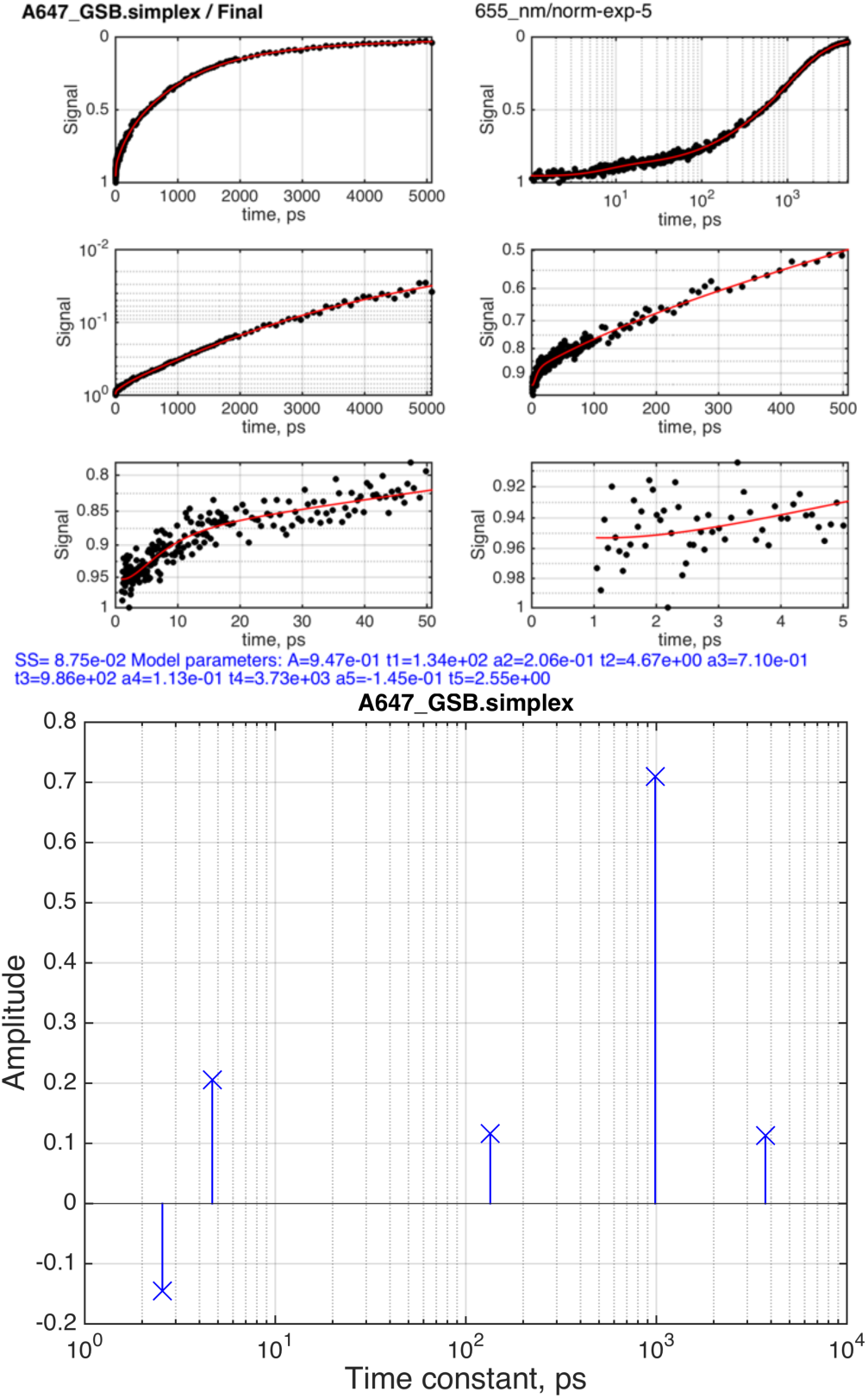
Five-exponential fit.

## Hypothesis testing

In general, it is often possible to reduce model deviations from the data but adding additional variable parameters. However, addition of fitting parameters reduces number of degrees of freedom in the datasets. Therefore, relative improvement in the sum of squares of the fit is always achieved at a cost of reduced number of degrees of freedom^*6, 7*^. This relative cost-benefit effect is evaluated by *IDAP* using the corrected Akaike’s Information Criterion, AIC_c_, which is calculated as^*6, 8-10*^:

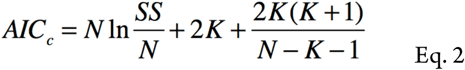

where N is the number of the data points, K—number of fitting parameter plus one, SS—sum of squares of residuals from the fit. The Evidence Ratio measures the relative likelihood of any pair of the models to accurately describe the data:

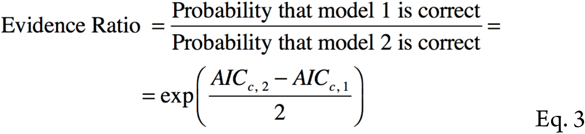

The *hypothesis_testing.m* script compares fitting results (stored in corresponding *session.mat* files) pairwise and produces the output in the form “Model X is more likely to be correct by the factor of Y”. Table 1 summarizes the testing results indicating that the four-exponential model is the one that optimally accounts for the shape of the decay curve. The five-exponential model is marginally better but includes a potentially meaningless term: negative amplitude with the sub-picosecond kinetics.

**Table 1.** Hypothesis testing to determine relative likelihood of the models to accurately describe the GSB decay of A647.

| Model 1 | Model 2 | Model 2 is more correct by the factor of, (1/ER) | Model 1 is more correct by the factor of, (ER) |
| --- | --- | --- | --- |
| single-exp | double-exp | 2e+93 |  |
| double-exp | triple-exp | 4e+31 |  |
| triple-exp | four-exp | 1e+03 |  |
| four-exp | five-exp | 4 |  |
| five-exp | six-exp |  | 200 |

The six-exponential model contains excessive number of parameters that makes fitting algorithm unstable that it cannot even reach the lowest sum-of-squares produced by the five-exponential model. This reminds us that fitting results are not the true answers but outcome of a “walk” of the optimization algorithm towards the global minimum. MATLAB includes fitting algorithms for improving convergence to the global minimum available in *IDAP* by setting the *fit_only_algorithm* parameter to ‘*DirectSearch’* or ‘*Global-Search’*. However, we did not observe better convergence with these complex (and very slow) algorithms, therefore, ‘*simplex’* remains a fitting method of choice for the TA data. Note, that simplex method ignores minimum and maximum values set for the parameters (as implemented in the MATLAB code).

## Confidence intervals of fitting parameters

The four-exponential model was found to be most likely to be correct. To determine confidential intervals and mutual correlation between parameters, we run *fit_exp_decay_main.m* with *fit_mode = ‘Monte-Carlo’* (which becomes the extension of the data-folder name). The algorithm, first, obtains the best-fit of the dataset. Then, the RMSD from the data is calculated and used to create mutliple simulated datasets perturbed with random noise. The model is sequentially fit to them accumulating distributions of best-fit values for the parameters. These distributions are percentiled at 2.5% and 97.5% of the population to extract 95% confidence interval. The confidence intervals are plotted on the graph for each subsequent simulation/fitting run for the user to follow the progress. As the population of best fit values becomes large enough the distribution’s 95% interval becomes less and less sensitive to subsequent addition of the new values. The *IDAP* algorithm monitors relative change of the confidence intervals as a criterion for stopping the simulations. The parameters controlling the simulations are listed in the code *TotalFit.m* in the section “Parameters controlling Monte-Carlo error estimates”. Figure 20 shows confidence intervals of amplitudes and lifetimes as rectangles defined by lower and upper boundaries of both parameters. Note, that confidence intervals are only shown for the N-1 amplitudes because *a*_*1*_ is a dependent parameter. It is reasonable to reassign confidence interval of the total amplitude *A* in Eq. 1 to the *a*_*1*_. (available from *adjustable_parameters_only.txt*).

**Figure 20.**
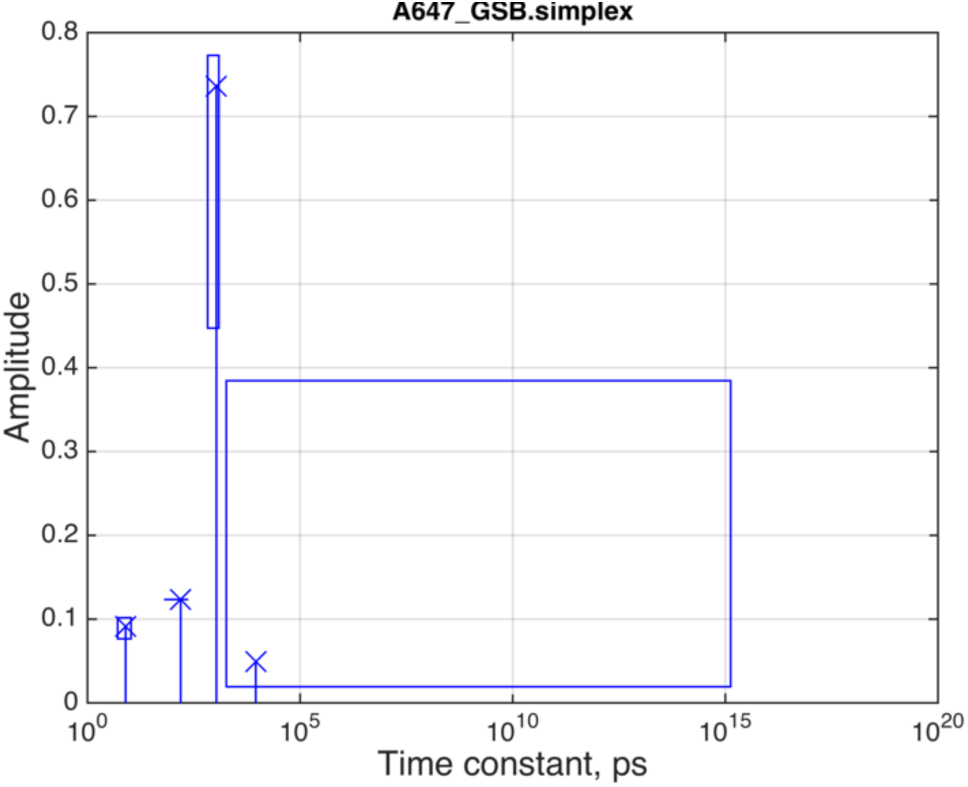
Four-exponential model 95% confidence intervals.

Figure 20 illustrates that the longest lifetime is extremely poorly defined. Even if it improves the fit over the three-exponential model, its value is, essentially, defined by just a minimum boundary of 1800 ps. It may be reasonable to return to the three-exponential model, which produces only marginally less perfect fit (5% greater sum of squares) yet yields more defined fitting parameters (Figure 21). Table 2 lists fitting parameters with their uncertainties for these two models. The *t*_*2*_ is similar in both models, while *t*_*1*_ and *t*_*3*_ are close. The difference of longer lifetimes between two models may be explained by the optimization routine attempting to “roll in” the deviations of the 4-5 ns time range into the longest lifetime of the triple-exponential model. In the four-exponential fit, that deviation is accounted for by the uncertain fourth lifetime and the t_2_ and t_3_ are only accounting for the shorter time-scale decay.

**Figure 21.**
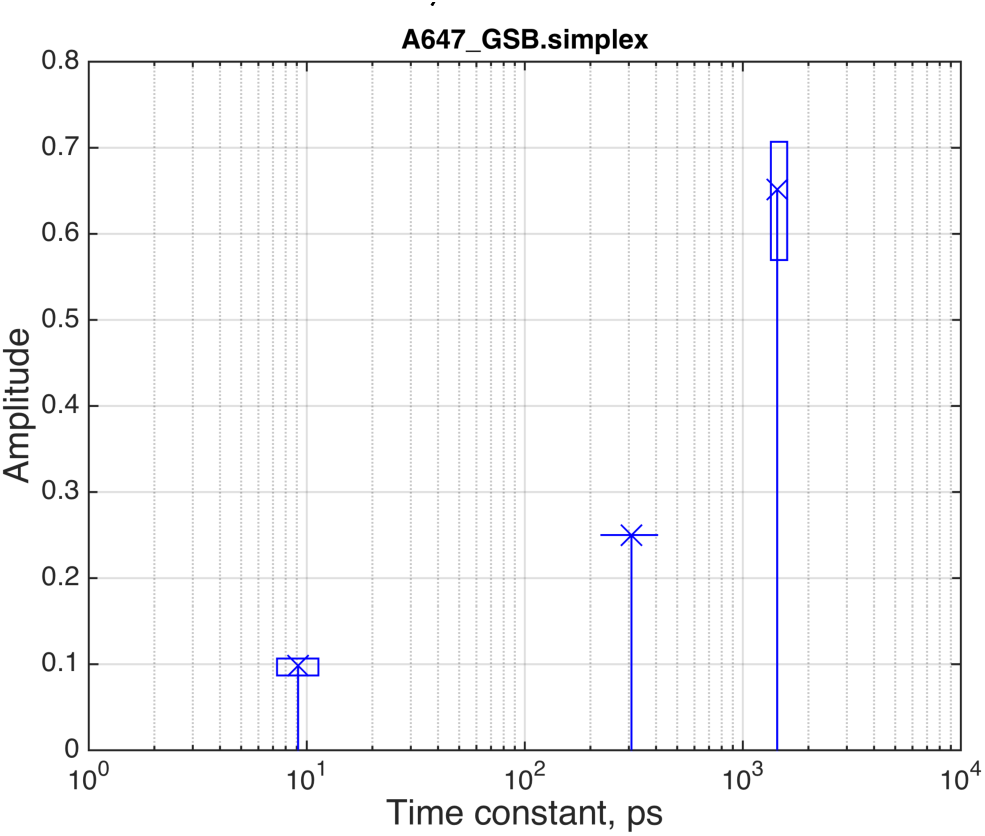
Three-exponential model 95% confidence intervals.

**Table 2.** Best fit parameters for the three- and four-exponential models.

| Parameter | best fit value | 95% confidence interval |
| --- | --- | --- |
| **Triple-exponential model** |  |  |
| $A$ | 0.97 | 0.96—0.98 |
| $a_2$ | 0.099 | 0.087—0.11 |
| $a_3$ | 0.65 | 0.57—0.71 |
| $t_1$ | 300 ps | 200—400 ps |
| $t_2$ | 9 ps | 7—11 ps |
| $t_3$ | 1400 ps | 1300—1600 ps |
| **Four-exponential model** |  |  |
| $A$ | 0.98 | 0.97—0.99 |
| $a_2$ | 0.09 | 0.08—0.10 |
| $a_3$ | 0.74 | 0.45—0.77 |
| $a_4$ | 0.05 | 0.02—0.4 |
| $t_1$ | 160 ps | 70—230 ps |
| $t_2$ | 8 ps | 5—10 ps |
| $t_3$ | 1100 ps | 700—1200 ps |
| $t_4$ | 9000 ps | >1800 ps |

Another observation from Table 2 is that the confidence intervals for the amplitudes in the four-state model are significantly increased. This is a signature that the model contains parameters not well supported by the data. As a result, the Monte-Carlo procedure yields broad confidence intervals. The lifetime of 9 ns might be a true feature of A647 or an artifact of the data collection but the broad confidence intervals prevent interpretation of this feature. As a conclusion, we prefer to go back to the three-state model as the most reasonable descriptor of the A647 GSB decay.

## Correlations between adjustable parameters

In the presence of noise in the data, the model parameters become less defined, which is reflected in increased confidence intervals. Correlation plots between parameters in Monte-Carlo runs show if the variation of one parameter may be compensated by adjustment of another. Figure 22 displays plots for parameters pairs of the triple-exponential model, where each blue point originates from one fitting run of a simulated noisy dataset. The black point indicates the best fit values of the original data. We observe that the amplitude and lifetime of the third exponential term demonstrates a significant degree of anticorrelation with the R value of 0.97, which means that reduction in the lifetime is compensated by increasing the amplitude. Therefore, both parameters cannot be determined more precisely with the existing data. High correlation also tells us that if one of the parameters was known from other data and fixed in the fit, the value of the second parameter could be very well defined. This will not be true for the first and second exponential terms, which parameters are virtually uncorrelated.

**Figure 22.**
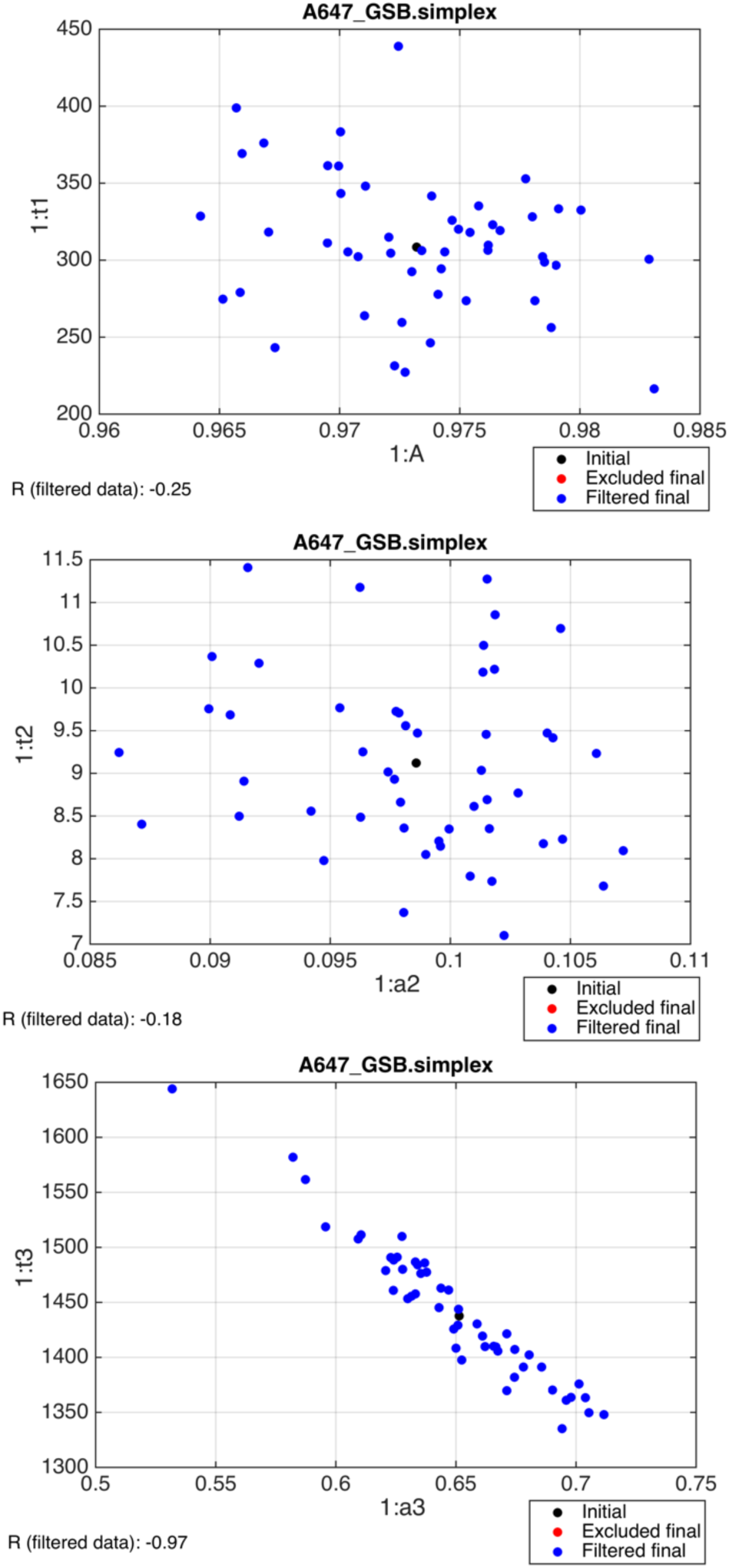
Correlation plots between A and t_1_ (top), a_2_ and t_2_ (middle) and a_3_ and t_3_ (bottom) parameters of the triple-exponential model.

## Summary

In this manuscript, I described the workflow for analysis of TA data using *IDAP* software including visualization of the spectral regions, extraction of the integrated decay traces, fitting and model selection based on Akaike’s Information Criterion.

## Supporting Information

Python TA data converter; MATLAB control scripts; plotting and hypothesis testing scripts.

## ACKNOWLEDGMENT

The author is thankful to Dr. Jier Huang and Brian Pattengale for recording the TA data and helpful discussions. The author acknowledges financial support of the Marquette University Committee on Research (2012 Summer Faculty Fellowship and 2012 Regular Research Grant).

